# Quercetin Mitigates Aluminium Chloride-Induced Neurotoxicity by Modulating Oxidative Stress, Neuroinflammation, and Neurotransmitter Dysregulation in Rats

**DOI:** 10.1101/2025.09.11.675655

**Authors:** Fatoki Tunji Micheal, Ajibola Toheeb Adesumbo, Onaolapo Olakunle James, Onaolapo Adejoke Yetunde

## Abstract

Memory impairment, characterized by reduced ability to recall facts, information, and experiences, is increasingly recognized as a major public health concern. This trend is largely driven by the rising prevalence of age-related cognitive decline and Alzheimer’s disease within the aging population. Aluminium chloride (AlCl□), a well-established neurotoxicant, induces neurobehavioral and biochemical alterations that mimic key features of neurodegenerative disorders, thus serving as a reliable experimental model for evaluating neuroprotective agents. This study assessed the neuroprotective efficacy of quercetin in a rat model of AlCl□-induced neurotoxicity. Fifty adult male rats (n = 10 per group) were randomly assigned into five groups. Neurotoxicity was induced by oral administration of AlCl□ (100 mg/kg/day) for 14 days. Subsequently, from days 14 to 35, rats received daily treatments of quercetin (100 or 200 mg/kg), donepezil (3 mg/kg), or vehicle control. AlCl□ exposure significantly impaired body weight gain, feed intake, locomotor activity, grooming behaviour, and spatial memory performance. Quercetin treatment markedly ameliorated these deficits, as evidenced by improved performance in Y-maze and radial-arm maze tasks. Biochemical analysis revealed that quercetin significantly reduced lipid peroxidation, enhanced total antioxidant capacity, and modulated inflammatory responses by decreasing pro-inflammatory cytokines (IL-1β, TNF-α) and elevating anti-inflammatory IL-10 levels. Furthermore, quercetin restored acetylcholine and brain-derived neurotrophic factor (BDNF) concentrations and preserved hippocampal cytoarchitecture, as demonstrated by histopathological assessment. These findings highlight quercetin’s therapeutic potential in mitigating aluminium-induced neurotoxicity and suggest its utility in the management of neurodegenerative disorders.

## 1. Introduction

Cognitive dysfunction and memory impairment represent early and prominent clinical manifestations of neurodegenerative disorders, commonly linked to structural and functional abnormalities within key brain regions such as the hippocampus and cerebral cortex [1-5]. Aluminum chloride (AlCl□), a well-characterized environmental neurotoxicant, is widely employed in experimental models to simulate neurodegenerative processes, owing to its capacity to disrupt neuronal architecture and impair cognitive functions [6].

There is ample evidence that AlCl□ exposure induces neuronal degeneration, characterised by cell loss, nuclear vacuolation, and histopathological damage in various brain regions, particularly the hippocampus [7]. Additionally, AlCl□ elevates oxidative stress markers, impairs cholinergic neurotransmission, and increases acetylcholinesterase (AChE) activity; factors that collectively contribute to synaptic dysfunction and cognitive deficits [1, 8–1]. There have also been reports that aluminium ions (Al^3^□) can mimic ferric ions (Fe^3^□) by binding to transferrin, facilitating their transport across the blood–brain barrier and leading to their accumulation in the brain [11]. This accumulation further exacerbates oxidative stress and neuroinflammation, both of which are implicated in the pathogenesis of Alzheimer’s disease and related neurodegenerative disorders [12, 13]. Experimental studies have demonstrated that aluminum chloride (AlCl□) impairs hippocampal function by disrupting the balance between excitatory and inhibitory neurotransmitters, interfering with synaptic signaling pathways [14, 15], and downregulating the expression of essential synaptic protein; alterations that are closely linked to deficits in learning and memory [16-18]. Both preclinical models and epidemiological studies in humans have consistently associated aluminum exposure with impairments in spatial memory and performance on hippocampus-dependent cognitive tasks [19].

Given the limitations of current pharmacological interventions, recent research has increasingly focused on naturally occurring compounds with neuroprotective potential [20, 21]. Quercetin, a dietary flavonoid widely present in fruits and vegetables, has garnered attention for its potent antioxidant and anti-inflammatory properties, which may counteract AlCl□-induced neurotoxicity [22, 23]. Notably, quercetin exhibits selective cytotoxicity, promoting apoptosis in damaged neurons while sparing healthy cells. This effect is mediated through the modulation of cell cycle regulators, including p21, phosphorylated retinoblastoma protein (pRb), cyclin B1, and cyclin-dependent kinase 1 (CDK1), thereby supporting neuronal viability and attenuating neurodegenerative processes [24]. Moreover, studies have demonstrated that quercetin preferentially localizes to mitochondria, which is a critical site of oxidative metabolism and vulnerability in neurodegenerative conditions; thereby enhancing its capacity to mitigate mitochondrial dysfunction [25]. Among the key mitochondrial defense systems is paraoxonase 2 (PON2), an intracellular antioxidant enzyme highly expressed in dopaminergic brain regions [26, 27]. The enzyme PON2 has been reported to play a crucial role in scavenging mitochondrial superoxide, and its deficiency has been associated with impaired mitochondrial bioenergetics and heightened susceptibility to oxidative damage [28, 29]. The co-localization of quercetin and PON2 within mitochondria would suggest a potential synergistic mechanism underlying their neuroprotective effects against oxidative stress-induced neuronal injury. Taken together, these findings would underscore the therapeutic potential of quercetin as a neuroprotective agent capable of mitigating aluminum-induced hippocampal injury.

## 2. Materials and Methodolo

### 2.1 Chemicals and Drugs

Quercetin capsules (500 mg; MRM Nutrition, USA), donepezil hydrochloride (10 mg; Sigma-Aldrich, USA), and aluminium chloride (AlCl□; Sigma-Aldrich, USA) were procured for this study. Assay kits for malondialdehyde (MDA; E4601), interleukin-10 (IL-10), tumor necrosis factor-alpha (TNF-α; K1052), and interleukin-1β (IL-1β; E4818) were procured from BioVision Inc. (Milpitas, CA, USA). Neurotransmitter quantification kits for acetylcholine (ACh) and brain-derived neurotrophic factor (BDNF) were obtained from Abcam Biotechnology (Cambridge, United Kingdom).

### 2.2 Animals and Ethical Approval

Adult male Wistar rats were obtained from Empire Breeders, Orioke-Ara, Osun State, Nigeria. Animals were individually housed in well-ventilated wooden cages under controlled environmental conditions (temperature: 25□±□2.5°C; relative humidity: 50–60%) and maintained on a 12-hour light/dark cycle (lights on at 07:00 h and off at 19:00 h). Rats had ad libitum access to standard rat chow and tap water throughout the duration of the experiment. All experimental procedures were conducted in accordance with the ethical standards for animal experimentation and were approved by the Ethics Review Committee of the Faculty of Basic Medical Sciences, Ladoke Akintola University of Technology (Approval No. ERC/FBMS/089/2025). The study also conformed to the European Council Directive 2010/63/EU for the protection of animals used for scientific purposes.

### 2.3 Diet and Supplementation

Rats were fed a standard rodent diet (Top Feeds®, Nigeria) formulated to contain 29% protein, 58% carbohydrates, and 11% fat per kilogram of feed. Throughout the experimental period, animals were assigned to receive either the standard diet alone or a diet supplemented with quercetin at concentrations of 100 mg/kg or 200 mg/kg of feed, as previously described [30].

### 2.4 Experimental Methods

Fifty adult male Wistar rats (150–170 g) were randomly divided into five experimental groups (n =10). Group A served as the normal control and received distilled water (vehicle) orally at a dose of 10 mL/kg body weight. Group B (AlCl□ control) was administered aluminum chloride (AlCl□) orally at 100 mg/kg/day. Groups C and D received quercetin at 100 mg/kg and 200 mg/kg body weight [30], respectively, in addition to AlCl□. Group E received donepezil at 3 mg/kg body weight alongside AlCl□ administration. AlCl□ was administered once daily from Day 1 to Day 14 to induce neurotoxicity. Quercetin and donepezil treatments were initiated on Day 14 and continued daily until Day 35. At the end of the treatment period (day 35), all animals underwent behavioural assessments, including the open field test, Y-maze, and radial-arm maze, to evaluate locomotor activity, exploratory behavior, and spatial working memory. Twenty-four hours after the final behavioural test, five rats from each group were randomly selected and euthanized via cervical dislocation. Blood samples were collected via intracardiac puncture for the analysis of systemic biochemical parameters, including pro-inflammatory cytokines (tumor necrosis factor-α [TNF-α], interleukin-1β [IL-1β]) and the anti-inflammatory cytokine interleukin-10 (IL-10), as well as lipid peroxidation (malondialdehyde [MDA]) and total antioxidant capacity (TAC). Brains were rapidly excised, visually inspected for gross abnormalities, and weighed. The hippocampus was dissected and either homogenized for biochemical analyses or fixed in formalin, paraffin-embedded, sectioned at 5 µm, and stained for histopathological evaluation. Supernatants from hippocampal homogenates were analyzed for acetylcholine (ACh) and brain-derived neurotrophic factor (BDNF).

### 2.5 Measurement of Relative Changes in Body Weight and Feed Intake

Body weight was recorded weekly, while daily feed intake was monitored using an electronic weighing balance (Mettler, Toledo, Type BD6000, Greifensee, Switzerland). The relative change in body weight and feed intake for each rat was calculated using the following equation, and the results were then analysed to determine the statistical mean for all animals.

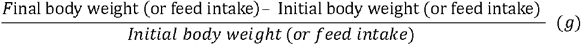

### 2.6 Behavioural Testing Procedure

Each rat was subjected to the behavioural test paradigms only once to avoid learning or habituation effects. Behavioural assessments were conducted in a quiet, controlled environment between 07:00 and 14:00 hours. Testing followed a standardized sequence: (1) Y-maze for spatial working memory, (2) open field test for locomotor and exploratory behaviour, and (3) radial-arm maze for spatial memory evaluation. On test days, animals were transported to the behavioural laboratory in their home cages and allowed to acclimate for approximately 30 minutes before testing. For each behavioural assay, the rat was gently placed in the respective apparatus, and its behaviour was recorded for subsequent analysis. Upon completion of each test, the rat was returned to its home cage. To eliminate potential olfactory cues and ensure consistent testing conditions, all apparatuses were thoroughly cleaned with 70% ethanol and allowed to dry between trials as previously described [31].

#### 2.6.1 Open field Novelty induced Behaviours

The open-field test was employed to evaluate general locomotor activity and central excitatory or inhibitory behaviors in rats, with a particular focus on self-grooming as an index of stereotypy. Each rat was placed in the open-field arena and observed for 10 minutes to assess horizontal locomotion, vertical exploration, and self-grooming behavior. The open-field apparatus consists of a rectangular wooden box measuring 72 × 72 × 26 cm, with the floor painted white and subdivided into 16 equal squares using permanent red lines. Behavioural scoring was conducted according to the procedure described by [32]. During the test, parameters recorded included horizontal locomotion or line crossing, quantified as the number of squares crossed, vertical locomotion measured by rearing frequency (number of times the rat stood upright on its hind limbs or contacted the walls) and self-grooming episodes counted as the number of discrete grooming bouts, including body cleaning, facial washing, and genital grooming. All behavioural data were recorded manually or with video assistance for subsequent analysis. The arena was thoroughly cleaned with 70% ethanol between trials to eliminate residual olfactory cues.

#### 2.6.2 Y-Maze and Radial-Arm Maze Tests

The Y-maze and radial-arm maze tests are employed to evaluate spatial working memory, a critical cognitive function necessary for exploration and adaptive behaviour. In rodents, spatial working memory is commonly assessed through spontaneous alternation behaviour, defined as the innate tendency to explore new arms of a maze without external reinforcement. This behavior reflects the animal’s ability to recall previously visited locations and is typically measured over a brief exploration period [32]. The Y-maze used in this study consisted of three arms arranged at 120° angles, each arm measuring 50 cm in length, 20 cm in height, and 10 cm in width. Each rat was placed at the end of one arm and allowed to explore freely for 5 minutes. An entry was counted when all four limbs had entered an arm. The sequence of arm entries was recorded to calculate the percentage of spontaneous alternation, defined as the number of triads containing entries into all three arms (i.e., ABC, BCA, or CAB) divided by the total possible alternations: The percentage of alternation was calculated as {(Actual alternations/Total arm entries – 2) x 100} over a 5-minute period, as outlined by [33]. This metric reflects the rat’s ability to remember and alternate between different arms of the maze, indicating intact spatial working memory. To further assess working memory performance, the eight-arm radial maze was used. The apparatus comprised eight arms (33 cm in length) extending from a central platform, equidistantly spaced in a radial pattern. During testing, each rat was placed on the central platform and allowed to explore freely for a set time period. The task required the animal to visit each arm without repeating entries. A working memory error was defined as a re-entry into an arm that had already been visited within the same trial. The number of errors was recorded as a direct index of spatial memory deficits.

#### 2.6.2 Elevated plus maze

The elevated plus maze (EPM) was employed to assess anxiety-related behaviour in rats. The apparatus consisted of four arms arranged in a plus (+) configuration, elevated 50 cm above the floor. Two opposing arms were open (25 × 5 × 0.5 cm), and two were closed (25 × 5 × 16 cm), with a central platform measuring 5 × 5 × 0.5 cm. The closed arms were enclosed by opaque walls to prevent falls, while the open arms had no side walls. Following administration of the test compound or vehicle, each rat was placed on the central platform facing one of the closed arms. Behavior was recorded for a 5-minute period, as described in previous studies [33-35]. An arm entry was defined as the rat placing all four limbs into the arm. Between trials, the maze was thoroughly cleaned with 5% ethanol to eliminate olfactory cues. The percentage of time spent in the arms was calculated as time in open arms or closed arms/total time x100. These measures reflect exploratory behavior and anxiety-like responses, with increased open-arm exploration typically interpreted as reduced anxiety.

### 2.7 Homogenisation of hippocampal tissue

Within 24 hours following the completion of behavioural assessments, five rats from each experimental group were euthanized by cervical dislocation. Brains were immediately excised, and the hippocampus was carefully dissected on an ice-cold surface. Each hippocampal sample was homogenized in ice-cold phosphate-buffered saline (PBS; pH 7.4) at a ratio of 1:10 (w/v) using a Teflon-glass homogenizer to preserve protein integrity. The homogenates were subsequently centrifuged at 5,000 rpm for 15 minutes at 4□°C to remove nuclei and cellular debris. The resulting supernatants were carefully collected and stored at −20□°C until further biochemical analyses were conducted for the determination of oxidative stress markers, cytokines, and neurotransmitter levels.

### 2.8 Biochemical Test

#### 2.8.1 Lipid Peroxidation Assay

Lipid peroxidation was evaluated by quantifying malondialdehyde (MDA) levels using the thiobarbituric acid-reactive substances (TBARS) assay, as previously described [36]. In this assay, MDA reacts with thiobarbituric acid (TBA) under high temperature and acidic conditions to form a pink-colored MDA–TBA adduct. The resulting complex was measured spectrophotometrically at an absorbance of 532 nm. The concentration of MDA in the hippocampal supernatants was calculated using a standard curve and expressed in micromoles per liter (μmol/L), serving as an index of lipid peroxidation and oxidative stress.

#### 2.8.2. Total Antioxidant Capacity Assay

Total antioxidant capacity (TAC) was assessed using the Trolox Equivalent Antioxidant Capacity (TEAC) assay, which measures the ability of antioxidants in the sample to scavenge or inhibit oxidized products, thereby reflecting overall antioxidant potential. The assay was performed using a commercially available kit, following the manufacturer’s protocol (BioVision Inc., Milpitas, CA, USA). Results were expressed in terms of Trolox equivalents, based on a standard curve generated with known concentrations of Trolox, a water-soluble vitamin E analog commonly used as a reference antioxidant.

#### 2.8.3 Acetylcholine and Brain-Derived Neurotrophic Factor (BDNF) Assays

The concentrations of acetylcholine and brain-derived neurotrophic factor (BDNF) in hippocampal homogenate supernatants were quantified using enzyme-linked immunosorbent assay (ELISA) kits, following the manufacturer’s protocols (Abcam Biotechnology, Cambridge, UK). Assays were conducted in accordance with the kit instructions, and absorbance was measured using a microplate reader. Final concentrations were calculated from standard curves and expressed in ng/mL for BDNF and μmol/L for acetylcholine, as appropriate

#### 2.8.4 Cytokine Quantification

Hippocampal homogenate levels of tumor necrosis factor-alpha (TNF-α), interleukin-10 (IL-10), and interleukin-1β (IL-1β) were determined using enzyme-linked immunosorbent assay (ELISA) kits, according to the manufacturer’s instructions (BioVision Inc., Milpitas, CA, USA). The assays quantified the total concentrations (both bound and unbound forms) of each cytokine. Absorbance readings were taken using a microplate reader, and cytokine concentrations were calculated based on standard curves and expressed in pg/mL [37].

### 2.9 Histological Processing and Staining

Following completion of behavioural assessments, rat brains were carefully dissected and grossly examined for visible abnormalities. Hippocampal sections were fixed in 10% neutral buffered formalin for 48 hours, then processed for routine paraffin embedding. Tissue blocks were sectioned at a thickness of 5□µm using a rotary microtome. Sections were stained with hematoxylin and eosin (H&E) for general histoarchitecture, Cresyl Fast Violet for Nissl substance visualization, and silver stain to assess neuronal fiber integrity. All histological procedures were performed according to the methods described by [38].

### 2.10 Histopathological Examination

Histologically processed hippocampal sections were examined using a Sellon-Olympus trinocular light microscope (Model XSZ-107E, China) equipped with a Canon PowerShot A2500 digital camera for photomicrography. Representative photomicrographs were captured for qualitative histopathological evaluation. All analyses were performed by an experienced pathologist blinded to the treatment groups to ensure unbiased assessment of tissue morphology.

### 2.11 Statistical Analysis

Data were analysed using **Chris Rorden’s ANOVA for Windows** (version 0.98). Group comparisons were performed using one-way analysis of variance (ANOVA), followed by **Tukey’s Honestly Significant Difference (HSD)** post-hoc test for multiple comparisons. All results are presented as **mean ± standard error of the mean (SEM)**. A p-value of **< 0.05** was considered statistically significant.

## 3. Results

### 3.1 Effect of quercetin on body weight

Figure 1 shows the effect of **quercetin** on the relative change in **body weight** following **aluminium chloride (AlCl**O**)-induced neurotoxicity** in rats. A significant reduction in body weight gain **(p < 0.001)** was observed in the **AlCl**□, **AlCl**□**/Q100, AlCl**□**/Q200**, and **AlCl**□**/Done** groups compared to the **control group**. However, when compared to the **AlCl**□ **group**, body weight gain was significantly higher in the **AlCl**□**/Q100, AlCl**□**/Q200**, and **AlCl**□**/Done** groups. Additionally, body weight gain in the **AlCl**□**/Q100** and **AlCl**□**/Q200** groups was significantly higher than in the **AlCl**□**/Done** group.

**Figure 1.**
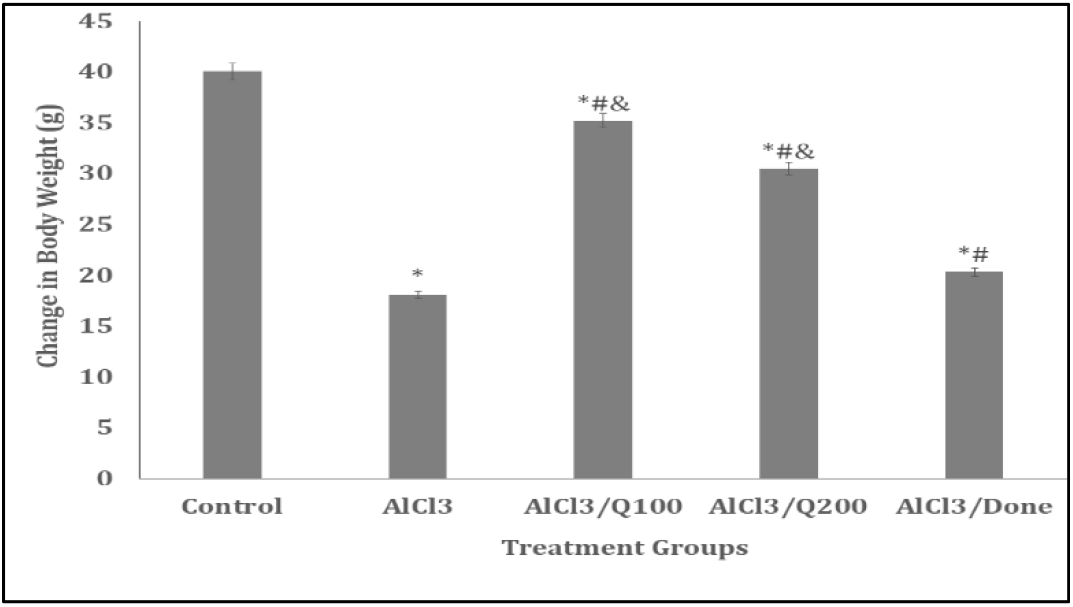
Effect of quercetin on body weight change. Data presented as Mean ± S.E.M, *p<0.05 significant difference from control. #p<0.05 significant difference from AlCl□, ^&^p<0.05 significant difference from AlCl□/Done. Number of rats per treatment group =10. AlCl□: Aluminium chloride, Done: Donepezil, Q: Quercetin.

### 3.2 Effect of quercetin on feed intake

Figure 2 shows the effect of quercetin on the relative change in feed intake following aluminium chloride (AlCl□)-induced neurotoxicity in rats. A significant reduction in feed intake (p < 0.001) was observed in the AlCl□, AlCl□/Q100, AlCl□/Q200, and AlCl□/Done groups compared to the control group. However, relative to the AlCl□ group, feed intake was significantly higher in the AlCl□/Q100, AlCl□/Q200, and AlCl□/Done groups. Furthermore, feed intake in the AlCl□/Q100 and AlCl□/Q200 groups was significantly higher than that in the AlCl□/Done group.

**Figure 2.**
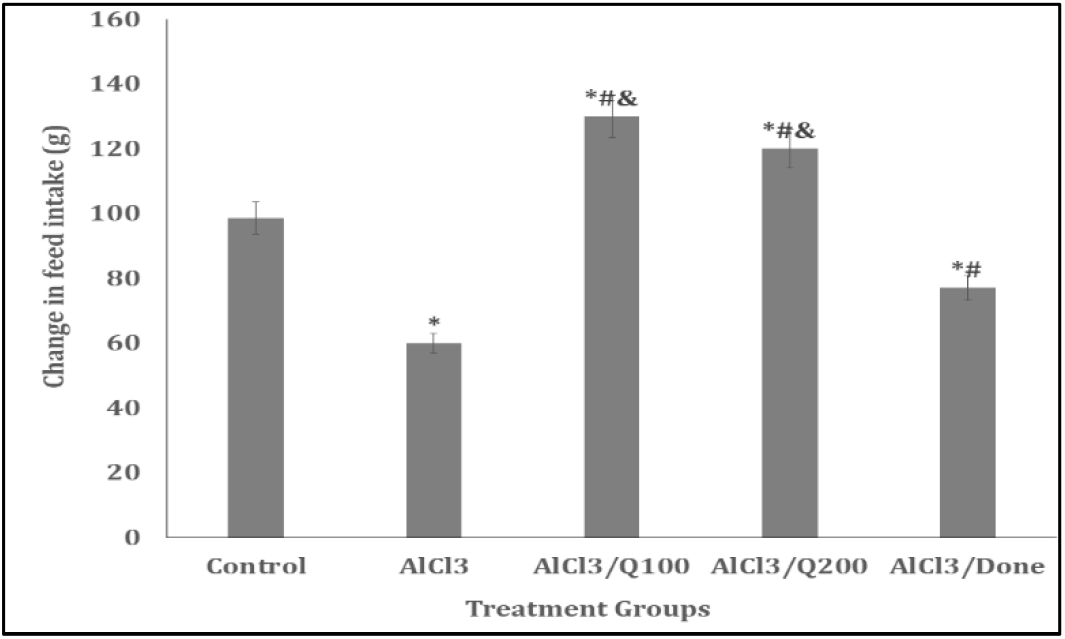
Effect of quercetin on feed intake. Data e presented as Mean ± S.E.M, *p<0.05 significant difference from control. #p<0.05 significant difference from AlCl□, ^&^p<0.05 significant difference from AlCl□/Done. Number of rats per treatment group =10. AlCl□: Aluminium chloride, Done: Donepezil, Q: Quercetin.

### 3.3 Effect of quercetin on horizontal locomotion

Figure 3 shows the effect of **quercetin** on **horizontal locomotion**, measured by the number of lines crossed in the **open field arena**, following **AlCl**□**-induced neurotoxicity** in rats. A significant decrease in locomotor activity **(p < 0.001)** was observed in the **AlCl**□ and **AlCl**□**/Done** groups compared to the **control**. In contrast, the **AlCl**□**/Q100** and **AlCl**□**/Q200** groups exhibited a significant increase in locomotor activity relative to the **control**. Furthermore, when compared to the **AlCl**□ **group**, locomotor activity was significantly higher in both the **AlCl**□**/Q100** and **AlCl**□**/Q200** groups. Similarly, locomotor activity was significantly increased in these groups compared to the **AlCl**□**/Done** group.

**Figure 3.**
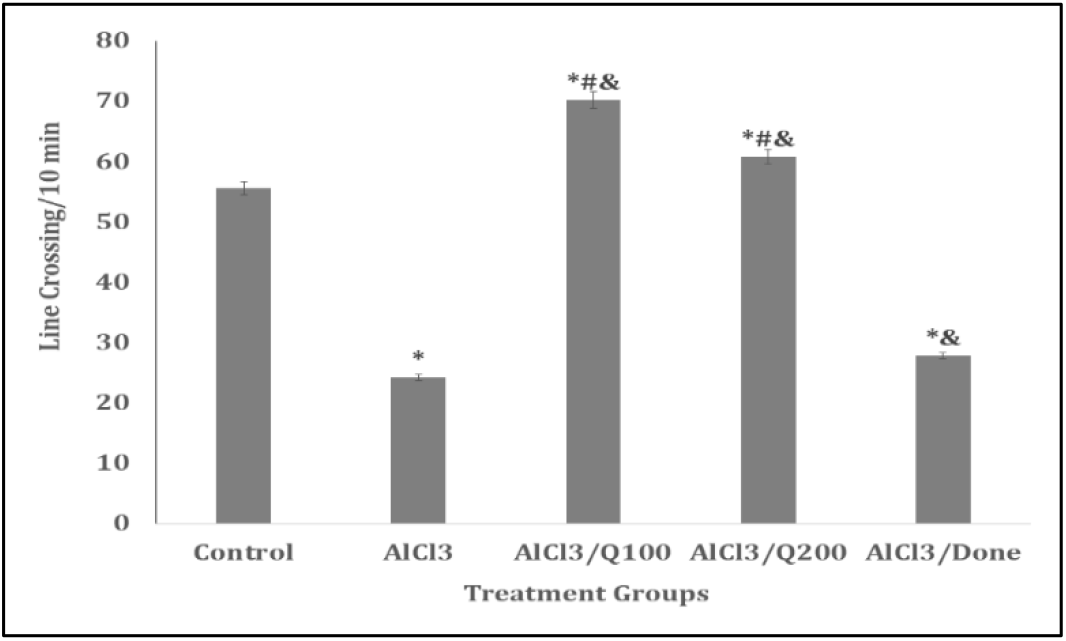
Effect of quercetin on horizontal locomotion. Data presented as Mean ± S.E.M, *p<0.05 significant difference from control. #p<0.05 significant difference from AlCl□, ^&^p<0.05 significant difference from AlCl□/Done. Number of rats per treatment group =10. AlCl□: Aluminium chloride, Done: Donepezil, Q: Quercetin.

### 3.4 Effect of quercetin on vertical locomotion

Figure 4 shows the effect of **quercetin** on **vertical locomotion**, measured by the number of **rears** in the **open field arena**, following **AlCl**□**-induced neurotoxicity** in rats. A significant reduction in rearing activity **(p < 0.001)** was observed in the **AlCl**□ and **AlCl**□**/Done** groups compared to the **control**. In contrast, the **AlCl**□**/Q100** and **AlCl**□**/Q200** groups showed a significant increase in rearing activity relative to the **control**. When compared to the **AlCl**□ **group**, rearing activity was significantly elevated in the **AlCl**□**/Q100, AlCl**□**/Q200**, and **AlCl**□**/Done** groups. Additionally, rearing activity in the **AlCl**□**/Q100** and **AlCl**□**/Q200** groups was significantly higher than in the **AlCl**□**/Done** group.

**Figure 4.**
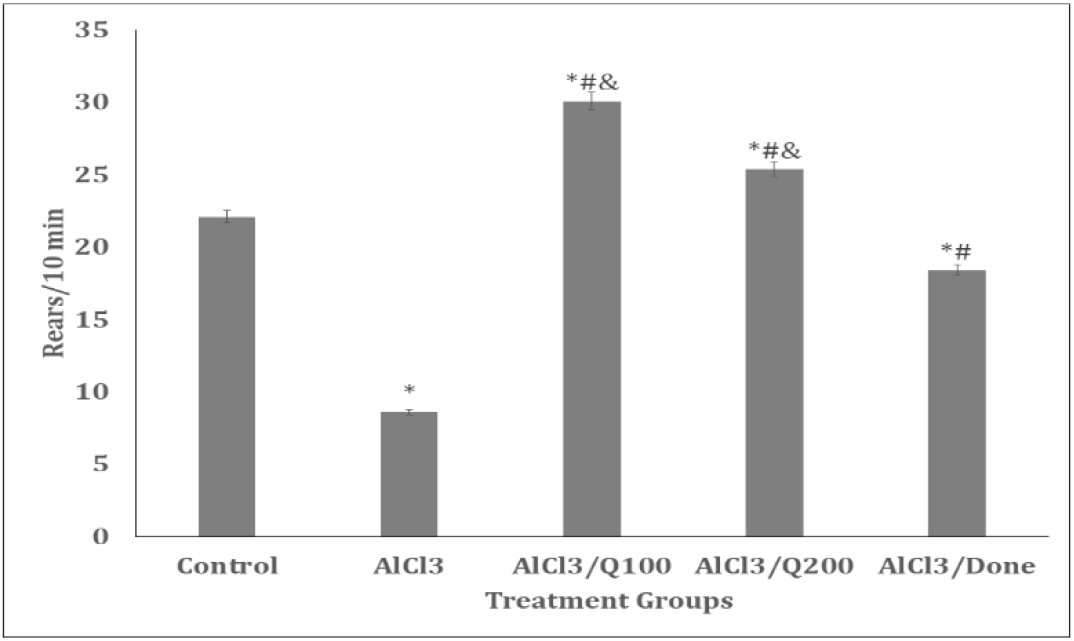
Effect of quercetin on vertical locomotion. Data presented as Mean ± S.E.M, *p<0.05 significant difference from control. #p<0.05 significant difference from AlCl□, ^&^p<0.05 significant difference from AlCl□/Done. Number of rats per treatment group =10. AlCl□: Aluminium chloride, Done: Donepezil, Q: Quercetin

### 3.5 Effect of quercetin on the self-grooming behaviours

Figure 5 shows the effect of quercetin on self-grooming behaviour in the open field arena following AlCl□-induced neurotoxicity in rats. A significant reduction in self-grooming (p < 0.001) was observed in the AlCl□, AlCl□/Q100, and AlCl□/Q200 groups compared to the control. However, relative to the AlCl□ group, self-grooming significantly increased in the AlCl□/Q100, AlCl□/Q200, and AlCl□/Done groups. Notably, when compared to the AlCl□/Done group, self-grooming was significantly reduced in both the AlCl□/Q100 and AlCl□/Q200 groups.

**Figure 5.**
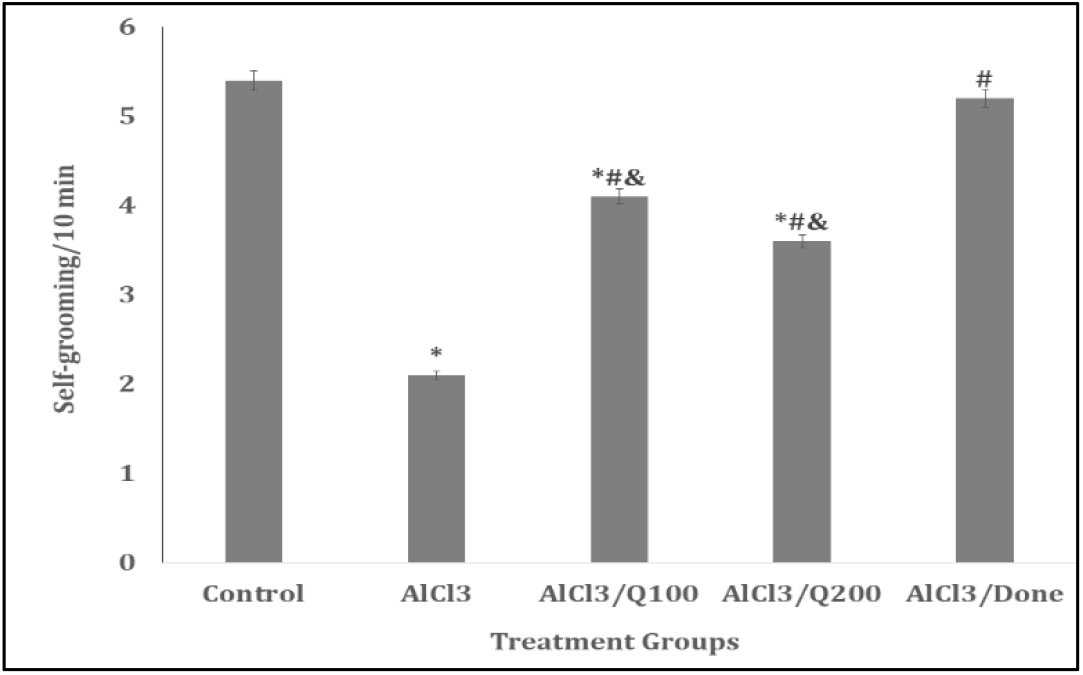
Effect of quercetin on self-grooming. Data presented as Mean ± S.E.M, *p<0.05 significant difference from control. #p<0.05 significant difference from AlCl□, ^&^p<0.05 significant difference from AlCl□/Done. Number of rats per treatment group =10. AlCl□: Aluminium chloride, Done: Donepezil, Q: Quercetin.

### 3.6 Effect of quercetin Y-maze spatial working memory

Figure 6 shows the effect of quercetin on spatial working memory, measured as the percentage of spontaneous alternation in a 5-minute Y-maze test following AlCl□-induced neurotoxicity in rats. A significant reduction (p < 0.001) in % alternation was observed in the AlCl□-treated group compared to the control, indicating marked impairment in spatial working memory. Co-administration of quercetin at both 100 mg/kg (AlCl□/Q100) and 200 mg/kg (AlCl□/Q200), as well as donepezil (AlCl□/Done), significantly improved % alternation compared to the AlCl□ group (p < 0.05 or as appropriate). Also, spatial working memory performance in the AlCl□/Q100 and AlCl□/Q200 groups was higher than that observed in the AlCl□/Done group, suggesting a potentially greater efficacy of quercetin in ameliorating AlCl□-induced cognitive deficits.

**Figure 6.**
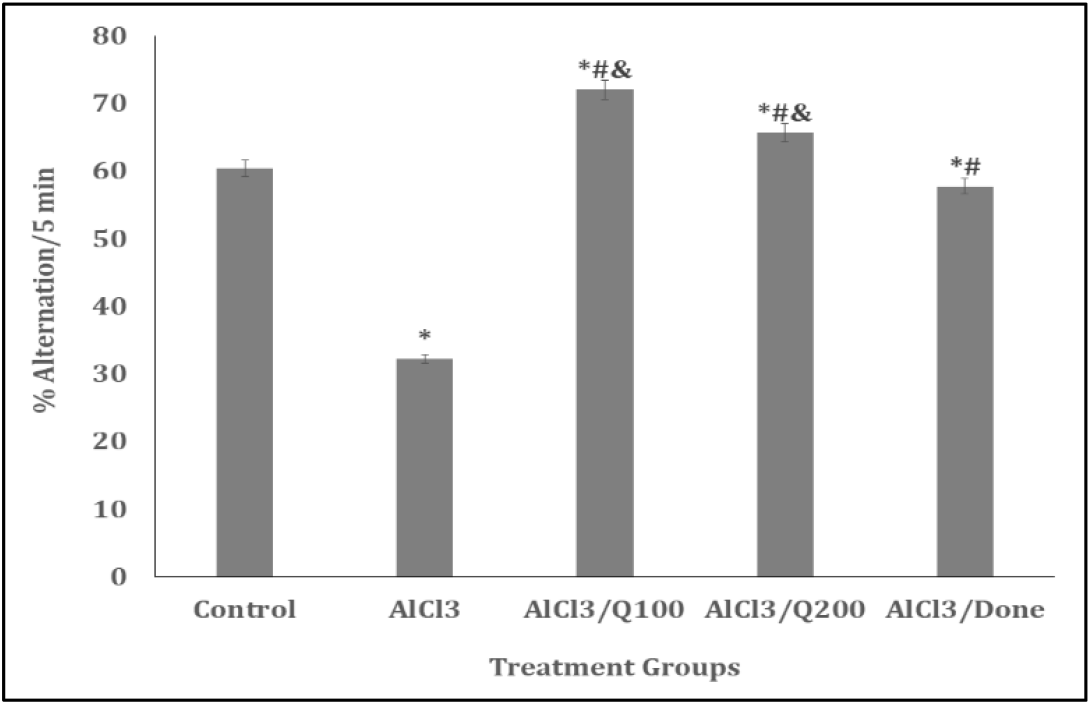
Effect of quercetin on Y-maze spatial working memory. Data presented as Mean ± S.E.M, *p<0.05 significant difference from control. #p<0.05 significant difference from AlCl□, ^&^p<0.05 significant difference from AlCl□/Done. Number of rats per treatment group =10. AlCl□: Aluminium chloride, Done: Donepezil, Q: Quercetin.

### 3.7 Effect of quercetin on radial-arm maze working memory

Figure 7 shows the effect of quercetin on spatial working memory, assessed using the radial-arm maze and expressed as the alternation index during a 5-minute testing period following AlCl□-induced neurotoxicity in rats. A significant reduction (p < 0.001) in alternation index was observed in the AlCl□, AlCl□/Q100, AlCl□/Q200, and AlCl□/Done groups compared to the control, indicating impaired spatial working memory. However, relative to the AlCl□ group, treatment with quercetin (AlCl□/Q100 and AlCl□/Q200) and donepezil (AlCl□/Done) resulted in significant improvements in alternation index (p < 0.05 or as appropriate). No significant differences were observed between the AlCl□/Q100, AlCl□/Q200, and AlCl□/Done groups, suggesting comparable efficacy of quercetin and donepezil in restoring spatial memory function.

### 3.7 Effect of Quercetin on Oxidative Stress Markers and Inflammatory Cytokines

Table 1. shows the effects of quercetin supplementation on oxidative stress parameters including malondialdehyde (MDA) and total antioxidant capacity (TAC), pro- and anti-inflammatory cytokines [interleukin-1β (IL-1β), interleukin-10 (IL-10), and tumor necrosis factor-alpha (TNF-α)], following AlCl□-induced neurotoxicity in rats. Administration of aluminium chloride resulted in a significant increase in MDA levels (p < 0.001) and a corresponding decrease in TAC compared to the control group, reflecting elevated oxidative stress. Similarly, levels of pro-inflammatory cytokines IL-1β and TNF-α were significantly elevated (p < 0.001), while the anti-inflammatory cytokine IL-10 was markedly reduced (p < 0.001), indicating enhanced neuroinflammatory response. Co-administration of quercetin at both 100 mg/kg (AlCl□/Q100) and 200 mg/kg (AlCl□/Q200) significantly ameliorated these alterations. Specifically, MDA, IL-1β, and TNF-α levels were significantly reduced, whereas TAC and IL-10 levels were significantly elevated relative to the AlCl□-only group (p < 0.05). Comparable effects were observed in the donepezil-treated group (AlCl□/Done).

**Table 1:**
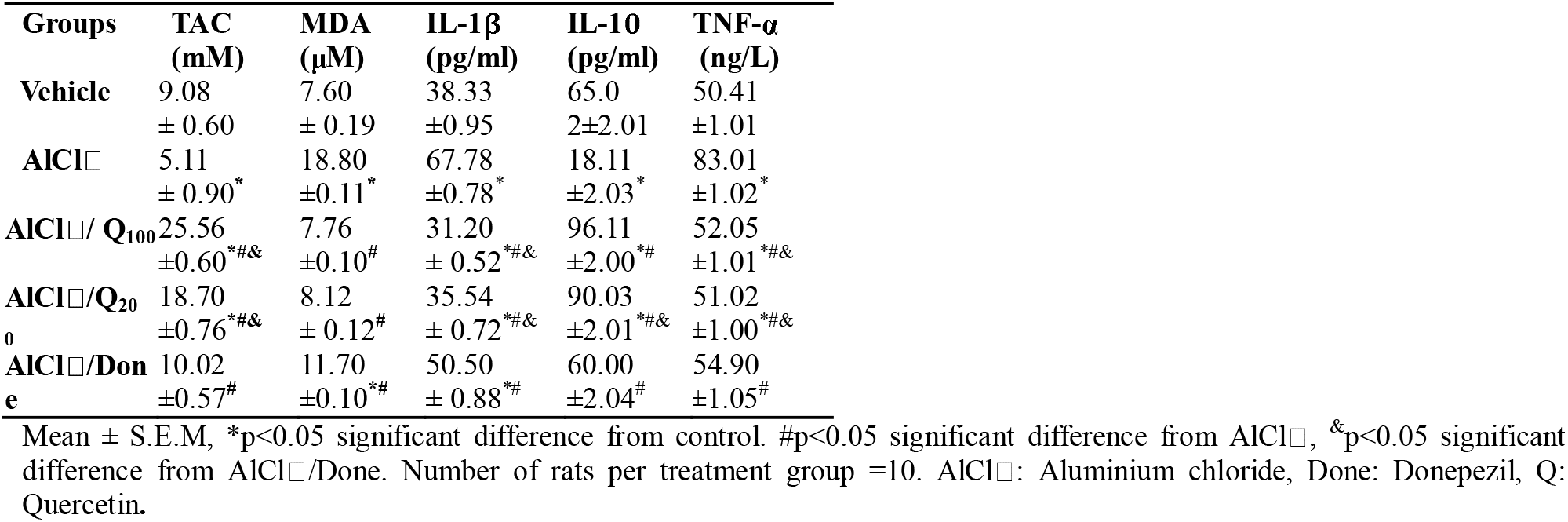
Effect of quercetin on oxidative stress and inflammatory parameters.

Importantly, quercetin at both doses produced significantly greater reductions in MDA and IL-1β levels, along with greater increases in TAC and IL-10 levels, compared to donepezil, suggesting superior antioxidant and anti-inflammatory efficacy. However, no significant differences were observed in TNF-α levels between the quercetin- and donepezil-treated groups.

### 3.8 Effect of Quercetin on hippocampal acetylcholine and brain derived neurotrophic factor levels

Table 2 shows the effect quercetin supplementation on neurotransmitter levels in the hippocampus. There was a significant decrease in acetylcholine levels with AlCl□ and an increase with AlCl□/Q100, AlCl□/Q200 and AlCl□/Done compared to control. Compared to AlCl□, acetylcholine levels increased with AlCl□/Q100, AlCl□/Q200 and AlCl□/Done, while compared to AlCl□/Done, a significant decrease in acetylcholine levels was observed with observed with AlCl□/Q200. Brain derived neurotrophic factor (BDNF) levels decreased significantly with AlCl□, AlCl□/Q200 and AlCl□/Done compared to control. Compared to AlCl□, BDNF levels increased with AlCl□/Q100, AlCl□/Q200 and AlCl□/Done, while compared to AlCl□/Done, BDNF levels increased with AlCl□/Q100.

**Table 2:** Effect of quercetin on hippocampal acetylcholine and brain derived neurotrophic factor levels.

| GROUPS | Acetylcholine<br>(pmol/mg) | BDNF<br>(pmol/mg) |
| --- | --- | --- |
| Vehicle | 4.2±0.15 | 25.10±0.30 |
| AlCl$_3$ | 1.34±0.12* | 10.01±0.50* |
| AlCl$_3$/Q200 | 6.51±0.13*#& | 27.45±0.60*#& |
| AlCl$_3$/Q200 | 5.62±0.13*#& | 20.20±0.44*#& |
| AlCl$_3$/Done | 2.55±0.12*# | 19.31±0.65*# |

Mean ± S.E.M, \*p<0.05 significant difference from control. #p<0.05 significant difference from AlCl $_3$ , &p<0.05 significant difference from AlCl $_3$ /Done. Number of rats per treatment group =10. AlCl $_3$ : Aluminium chloride, Done: Donepezil, Q: Quercetin, BDNF: Brain derived neurotrophic factor.

### 3.9 Effect of quercetin on hippocampal histomorphology

Figures 8 (a-e), 9 (a-e), 10 (a-e), 11 (a-e) and 12 (a-e) show representative photomicrographs of sections of the dentate gyrus region of the rat hippocampus stained using haematoxylin and eosin, Bielcholwsky’s silver stain and cresyl fast violet stains. Examination of the slides of animals in the control groups (8a, 9a, 10a, 11a and 12a) revealed triangular shaped layer of small granule cell neurons that make up the dentate gyrus. Also observed are numerous neuroglia and neuronal processes in the molecular layer that lie between the compact zones around the dentate gyrus. In the groups administered AlCl□ (8b, 9b, 10b, 11b and12b), graded loss of neurons and neuronal degeneration was observed. This is evidenced by poorly-staining nucleoli, loss of oval to round shape of the granule neuron, shrunken nucleus, and absence of the close-knit arrangement of the granule neurons. In the AlCl□/Q100 (8c, 9c,10c,11c and12c) and AlCl□/Q200 (8d, 9d, 10d, 11d and12d) groups, amelioration of AlCl□-induced changes in the dentate gyrus regions was observed. In the groups treated with AlCl□/Done (8e, 9e, 10e, 11e and 12e) although there was amelioration of the changes induced by AlCl□. However, the improvement observed with AlCl□/Q100 appeared to be better.

**Figure 8.**
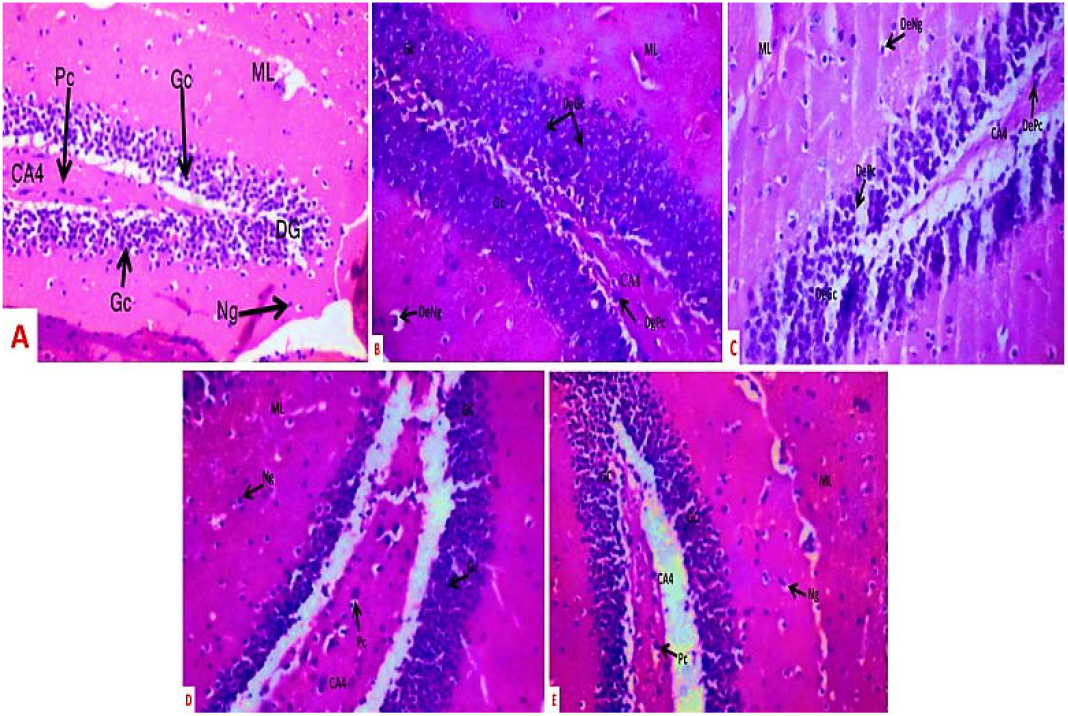
Representative photomicrographs of sections from the dentate gyrus region of the rat hippocampus stained with haematoxylin and eosin. Magnification: 100x. Gc: Granule cells, DePc: Degenerating pyramidal cells, Pc: Pyramidal cells.

**Figure 9.**
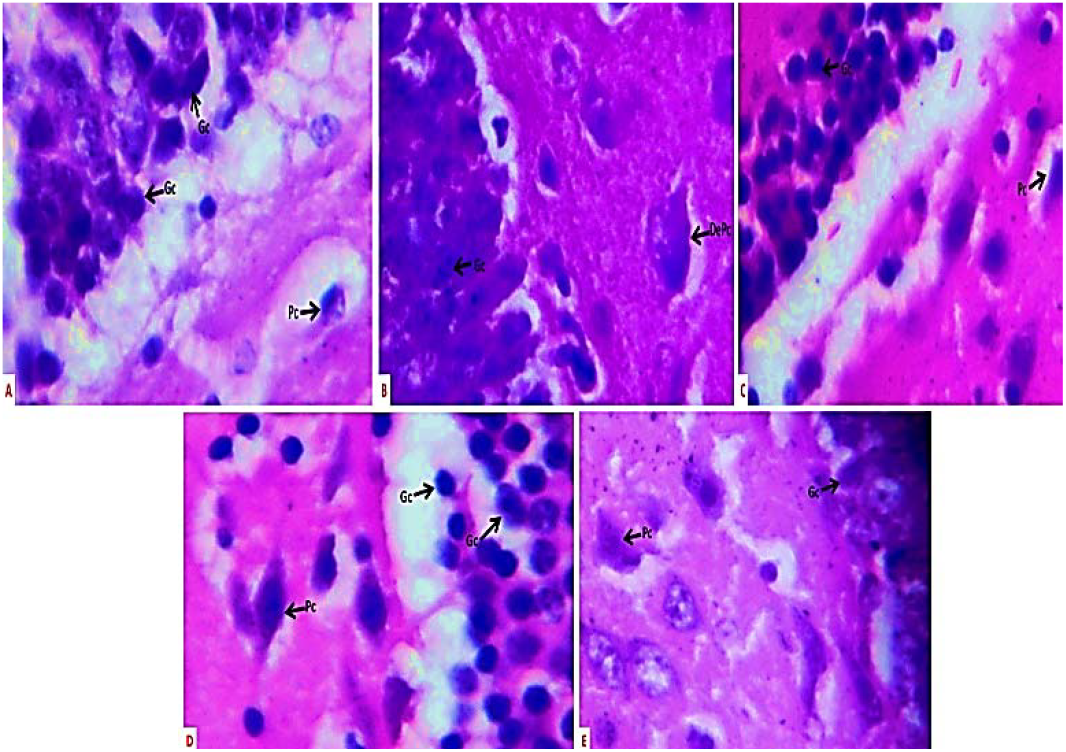
Representative photomicrographs of sections of the dentate gyrus region of the rat hippocampus stained using haematoxylin and eosin, magnification 400x. Gc: Granule cells, DePc: degenerating pyramidal cells, Pc: Pyramidal cells.

**Figure 10.**
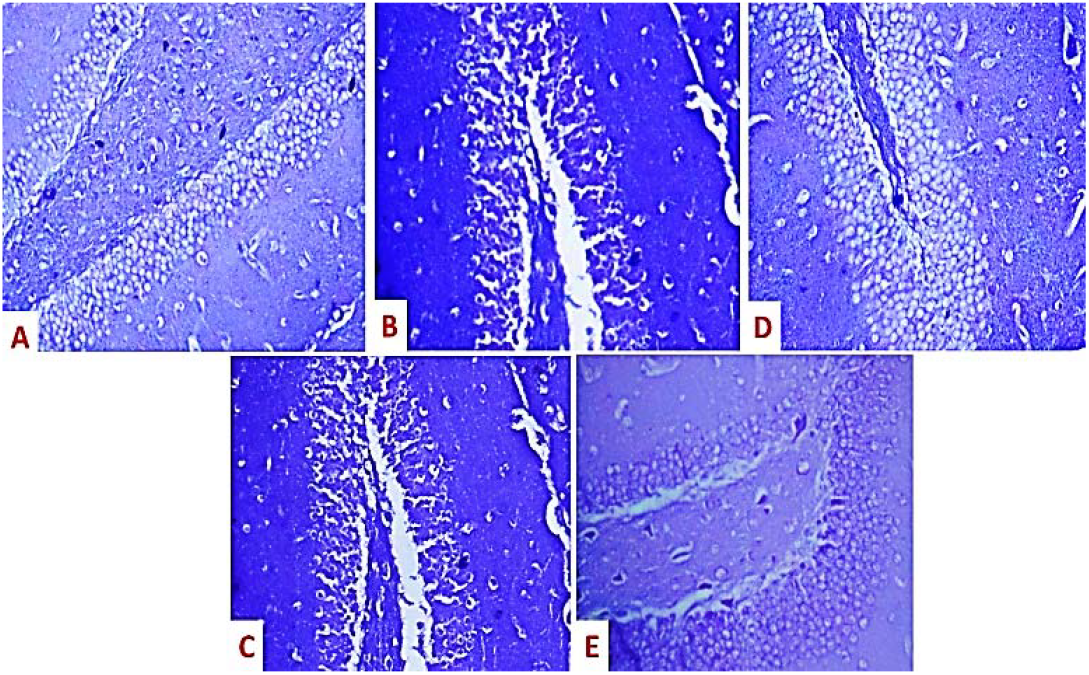
Representative photomicrographs of sections of the dentate gyrus region of the rat hippocampus stained using Cresyl fast violet stain. Magnification 100x

**Figure 11.**
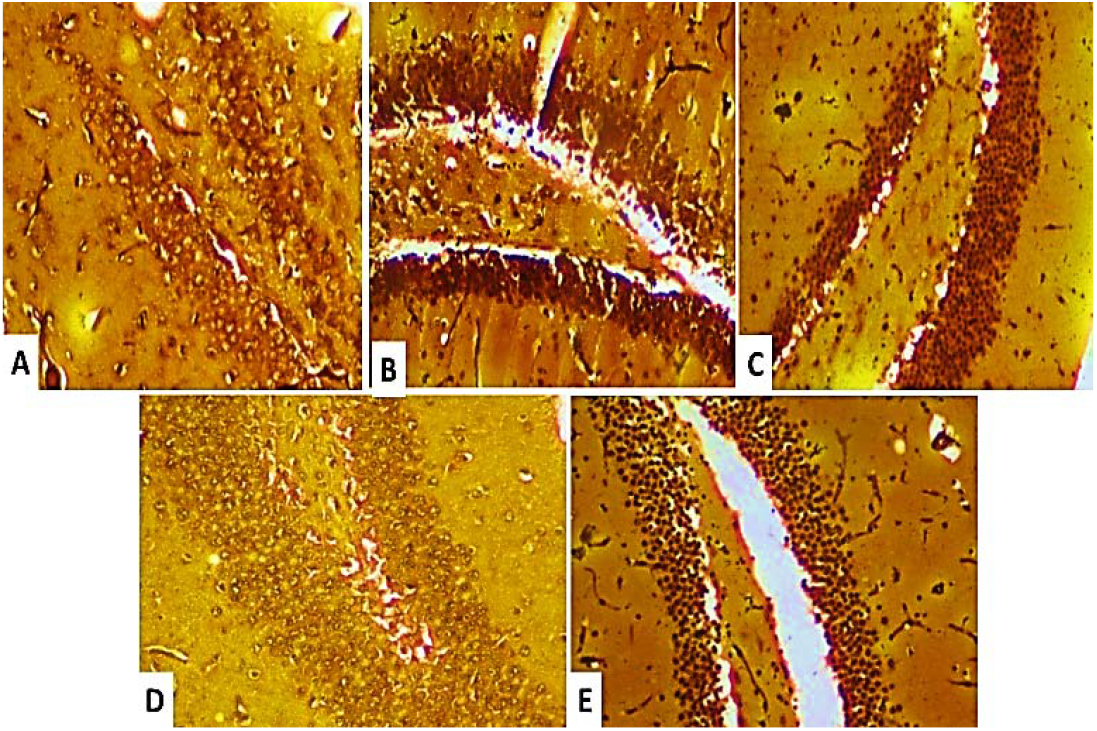
Representative photomicrographs of sections of the dentate gyrus region of the rat hippocampus stained using Bielschowsky silver stain. Magnification 100x

**Figure 12.**
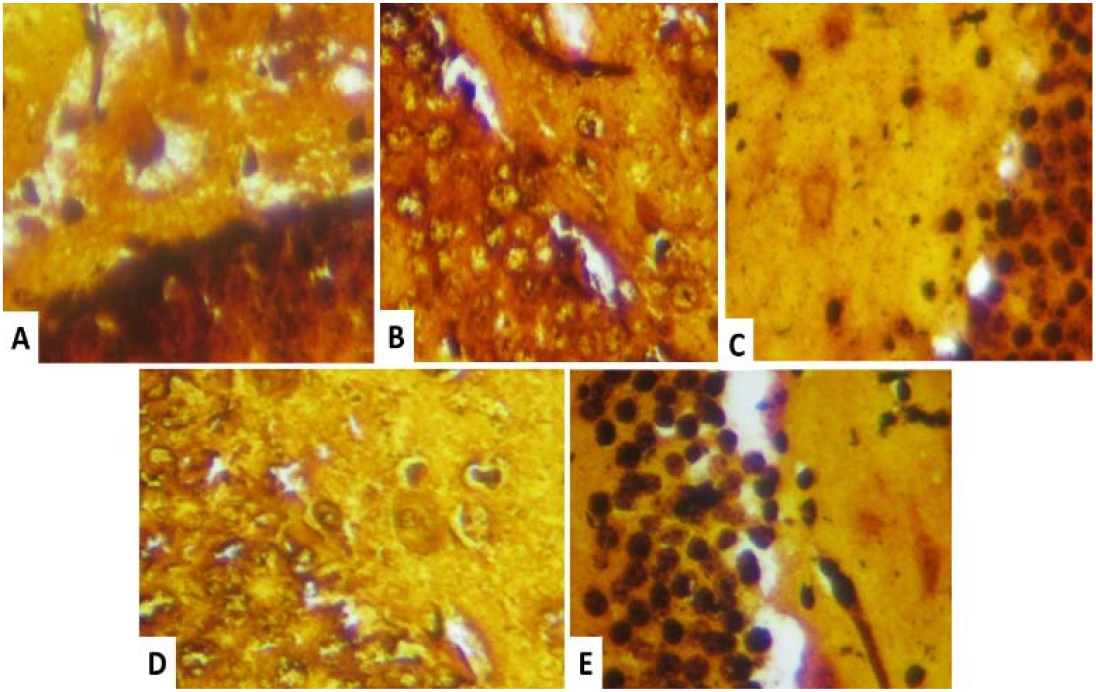
Representative photomicrographs of sections of the dentate gyrus region of the rat hippocampus stained using Bielschowsky silver stain. Magnification 400x

## 4. Discussion

Human exposure to aluminum is virtually unavoidable due to its widespread presence in metal recycling, ore processing, mining operations, processed foods, pharmaceuticals, and water purification systems [39]. The toxicological implications of AlCl□, especially its neurotoxic effects, have prompted increased interest in discovering compounds capable of mitigating its adverse effects [3, 4]. Quercetin, a naturally occurring flavonoid, has attracted significant attention for its antioxidant, anti-inflammatory, and neuroprotective properties [3,30]. The current study evaluated the efficacy of quercetin in alleviating AlCl□-induced neurotoxicity, focusing on behavioural alterations, oxidative stress parameters, inflammatory markers, neurotransmitter levels, and histological changes in the hippocampus.

The findings of this study revealed that AlCl□administration led to significant weight loss, consistent with previous reports [3, 4, 40]. This weight loss was associated with reduced feed intake, possibly due to a loss of appetite. Quercetin administration, especially at 100 mg/kg, significantly reversed the weight loss, corroborating earlier studies that had demonstrated its ability to modulate body weight [3, 30]. The observed restoration of feed intake further supports the role of quercetin in restoring energy balance.

Results of behavioural analyses revealed that AlCl□significantly impaired exploratory activities, as shown by decreased line-crossing and rearing frequencies in the open-field paradigm. These behaviors are indicators of motivation, motor activity, and anxiety. Quercetin administration significantly improved these behaviors, with the 100 mg/kg dose demonstrating effects comparable to/or better than those observed with donepezil. While grooming behaviour was also improved with quercetin treatment, the difference was not statistically significant compared to control, highlighting a possibly less sensitive response in this specific behaviour.

Y-maze testing demonstrated that AlCl□significantly impaired spatial working memory, evidenced by reduced spontaneous alternation behaviour. This supports existing literature linking aluminum exposure to cognitive deficits and dementia-like symptoms [3, 30]. Donepezil also showed memory enhancing effects corroborating previous reports and supporting its use as a nootropic agent [41]. Both doses of quercetin significantly improved spatial memory performance, with greater efficacy than donepezil, suggesting that quercetin may offer broader neuroprotective effects.

In agreement with prior findings [30, 42], this study also demonstrated that AlCl□ induces oxidative stress, evidenced by increased MDA and reduced TAC levels. Quercetin treatment reversed these changes, showing significant antioxidant activity. While donepezil also conferred some protective effects, quercetin, especially at 100 mg/kg produced superior outcomes. Similarly, the pro-inflammatory cytokines IL-6 and TNF-α were elevated following AlCl□ exposure and were significantly reduced with quercetin treatment, likely due to its modulation of intracellular pathways such as PI3K/Akt and MAPK/ERK [43, 44]. Neurochemical analysis revealed a significant reduction in hippocampal ACh levels in the AlCl□-treated group, consistent with impaired cholinergic signaling [45]. Quercetin restored ACh levels significantly, particularly at 100 mg/kg, and to a lesser extent at 200 mg/kg. Brain-derived neurotrophic factor levels were similarly suppressed by AlCl□, and quercetin reversed this suppression. The 100 mg/kg dose yielded greater BDNF restoration than the 200 mg/kg and donepezil-treated groups, indicating a dose-sensitive neurotrophic effect.

Histological analyses revealed marked neurodegeneration in the AlCl□-treated hippocampus, with loss of pyramidal neurons in the Cornu Ammonis region and granule cells in the dentate gyrus. Quercetin treatment markedly restored hippocampal architecture. Evidence from cresyl violet staining further supported this, showing improved neuronal integrity in the quercetin-treated groups compared to the poorly stained neurons in the AlCl□ group.

The hippocampus, known for its role in memory and spatial navigation, was particularly vulnerable to AlCl□-induced oxidative and inflammatory injury. The behavioural deficits observed in alternation index, line crossing, and rearing were thus consistent with the observed histopathology. Quercetin’s restorative effect on hippocampal structure and function underscores its neuroprotective potential. Results of hippocampal histomorphology following Bielcholwsky’s silver staining revealed the presence of neurofibrillary tangles, hallmark features of Alzheimer-like pathology, in AlCl□-treated rats. These dark brown intracytoplasmic inclusions were significantly reduced in quercetin-treated groups, with a greater reduction than seen in the donepezil group. NFTs, composed of hyperphosphorylated tau protein, contribute to synaptic dysfunction and cognitive impairment [46]. The reduction of NFTs with quercetin suggests its potential in modulating tau pathology.

## Conclusion

This study provides evidence that quercetin offers multifaceted neuroprotection against AlCl□-induced toxicity. It improves behavioural outcomes, restores oxidative and inflammatory balance, supports neurotransmitter and neurotrophic factor levels, and preserves hippocampal integrity. Importantly, quercetin at 100 mg/kg consistently outperformed the higher dose and donepezil in several parameters, suggesting an optimal therapeutic window. These findings support further investigation into quercetin as a potential adjunct or alternative therapy for aluminum-related neurotoxicity and neurodegenerative conditions.

## Availability of data and materials

Data generated during and analysed during the course of this study are available from the corresponding author on request.

## Declarations Ethics approval

Ethical approval for this study was granted by the Ethical Committee of the Faculty of Basic Medical Sciences (ERC/FBMS/089/2025).

## Competing interests

All authors of this paper declare that there is no conflict of interest related to the content of this manuscript.

Received 19th May 2025, Revised 23rd June 2025, Accepted 7th July, 2025, **Published online:25/08/25**

## References

1. Adelodun ST, Ishola OA, Abijo, A. Z., Olatunji, S. Y., Owolabi, J. O., Olanrewaju, J. A, Adekomi, D. A. Aluminium chloride-induced hippocampal damage: CA3 hippocampal subfield involvement and the neuroprotective role of Buchholzia coriacea ethanolic seed extract. Phytomedicine Plus, 2021;1(4), 100104.

2. Onaolapo AY, Obelawo AY, Onaolapo OJ. Brain Ageing, Cognition and Diet: A Review of the Emerging Roles of Food-Based Nootropics in Mitigating Age-related Memory Decline. Curr Aging Sci. 2019;12(1):2–14. doi: 10.2174/1874609812666190311160754.

3. Akinsehinwa AF, Samson PO, Onaolapo OJ, Onaolapo AY. Quercetin/Donepezil co-administration mitigates Aluminium chloride induced changes in open field novelty induced behaviours and cerebral cortex histomorphology in rats. Acta Bioscienctia 2025:1(1);01–07 10.71181/actabioscientia12110

4. Ajao JA, Akinsehinwa AS, Onaolapo OJ, Onaolapo AY. Alcohol Extract of Muira Puama (Ptychopetalum Olacoides) ameliorates Aluminium Chloride-induced changes in Behaviour and Cerebral cortex Histomorphology in Wistar Rats. Acta Bioscienctia 2025:1(1):022–029 10.71181/actabioscientia12140.

5. Onaolapo AY, Obelawo AY, Onaolapo OJ. Brain Ageing, Cognition and Diet: A Review of the Emerging Roles of Food-Based Nootropics in Mitigating Age-related Memory Decline. Curr Aging Sci. 2019;12(1):2–14. doi: 10.2174/1874609812666190311160754.

6. Rahman, A., Banu, Z. Impact of Aluminium Chloride (AlCl□) on Brain Function: A Review of Neurotoxic Mechanisms and Implications for Alzheimer’s Disease. Trends in Pharmaceutical Sciences, 2024; 10(4), 355–366.

7. Kumar A, Agrawal M, Murti Y, Behl S. Kumar S, Chaudhary H, Rastogi S. Safeguarding Neuronal Integrity: Unveiling Possible Role of NFκB in the Neuroprotective Efficacy of Andrographolide Contrary to Aluminium Chloride-induced Neurotoxicity and Associated Spatial Memory Impairments in Rats. Central Nervous System Agents in Medicinal Chemistry, 2025; 25(2), 157–168.

8. Firdaus, Z., Kumar, D., Singh, S. K., & Singh, TD Centella asiatica alleviates AlCl3-induced cognitive impairment, oxidative stress, and neurodegeneration by modulating cholinergic activity and oxidative burden in rat brain. Biological Trace Element Research, 2024; 200(12), 5115–5126.

9. Wen P, Sun Z, Gou F, Wang J, Fan Q, Zhao D, Yang L Oxidative Stress and Mitochondrial Impairment: Key Drivers in Neurodegenerative Disorders. Ageing Research Reviews, 2025; 102667.

10. Imam A, Oyegbola, C, Busari M, Gbemisola AM, Abubakar L, Odunayo AT, Ajao MS. λ-cyhalothrin induced sex-specific inflammation, glia activation and GABAergic interneuron disruption in the hippocampus of rats. BMC Pharmacology and Toxicology, 2025; 26(1), 22.

11. Walton J R. Cognitive deterioration and associated pathology induced by chronic low-level aluminum ingestion in a translational rat model provides an explanation of Alzheimer’s disease, tests for susceptibility and avenues for treatment. International Journal of Alzheimer’s Disease, 2012; 2012.

12. Bondy SC. The neurotoxicity of environmental aluminum is still an issue. Neurotoxicology, 2010; 31, 575–581.

13. Singh T, Goel RK. Neuroprotective effect of Allium cepa L. in aluminium chloride induced neurotoxicity. NeuroToxicology 2015; 49, 1–7.

14. Alqarni SS, Afzal M, Alharbi KS, Alenezi SK, Alsahli TG, Zaidi S, Altyar AE, Ghaboura N, Kazmi I, Mantargi MJS, Imam F. Rosiridin Protects Against Aluminum Chloride-Induced Memory Impairment via Modulation of BDNF/NFκB/PI3K/Akt Pathway in Rats. Medicina (Kaunas). 2024 Nov 4;60(11):1812. doi: 10.3390/medicina60111812.

15. Skalny AV, Aschner M, Jiang Y, Gluhcheva YG, Tizabi Y, Lobinski R, Tinkov AA. Molecular mechanisms of aluminum neurotoxicity: Update on adverse effects and therapeutic strategies. Adv Neurotoxicol. 2021;5:1–34. doi: 10.1016/bs.ant.2020.12.001.

16. Luo R, Shen XN, Fang DM Neuroprotective effects of hydrogen sulfide on chronic aluminium exposure-induced oxidative stress and apoptosis in hippocampus of mouse. Neurotoxicology 2018;8:27–39

17. Sharma D, Kannan K, Jain SK Aluminium chloride-induced oxidative stress, DNA damage and apoptosis in human liver hepatocellular carcinoma cell line HepG2. Mol Cell Biochem.; 2019; 451(1–2):197–209

18. Jafari A, Joneidi E, Roghani M. The Protective Effect of Berberine Against Memory Impairment Induced by Intrahippocampal Injection of Aluminium Chloride in Rats. Qom University of Medical Sciences Journal, 2022; 16(8), 640–653.

19. Prakash A, Kumar A. Effect of N-acetyl cysteine against aluminium-induced cognitive dysfunction and oxidative damage in rats. Basic Clin Pharmacol Toxicol. 2009 Aug;105(2):98–104. doi: 10.1111/j.1742-7843.2009.00404.x.

20. Onaolapo OJ, Olofinnade AT, Ojo FO, Onaolapo AY. Neuroinflammation and Oxidative Stress in Alzheimer’s Disease; Can Nutraceuticals and Functional Foods Come to the Rescue? Antiinflamm Antiallergy Agents Med Chem. 2022;21(2):75–89. doi: 10.2174/1871523021666220815151559.

21. Onaolapo OJ, Onaolapo AY, Olowe OA, Udoh MO, Udoh DO, Nathaniel TI. Melatonin and Melatonergic Influence on Neuronal Transcription Factors: Implications for the Development of Novel Therapies for Neurodegenerative Disorders. Curr Neuropharmacol. 2020;18(7):563–577. doi: 10.2174/1570159X18666191230114339.

22. Davis JM, Murphy EA, Carmichael MD. Effects of the dietary flavonoid quercetin upon performance and health. Curr Sports Med Rep. 2009 Jul-Aug;8(4):206–13. doi: 10.1249/JSR.0b013e3181ae8959.

23. Rauf A, Imran M, Khan IA, Gilani SA, Mehmood Z, Mubarak MS. Anticancer potential of quercetin: A comprehensive review. Phytotherapy Research, 2018;32(11), 2109–2130.

24. Jeong JH, An JY, Kwon YT, Rhee JG, Lee YJ Effects of low dose quercetin: Cancer cell-specific inhibition of cell cycle progression. J. Cell. Biochem, 2009; 106(1), 73–82

25. Fiorani M, Guidarelli A, Cantoni O Mitochondrial reactive oxygen species: the effects of mitochondrial ascorbic acid vs untargeted and mitochondria-targeted antioxidants. International Journal of Radiation Biology, 2021; 97(8), 1055–1062.

26. Draganov DI, La Du BN. Pharmacogenetics of paraoxonases: a brief review. Naunyn Schmiedebergs Arch Pharmacol. 2004;369(1):78–88. doi: 10.1007/s00210-003-0833

27. Marsillach, B. Mackness, M. Mackness, F. Riu, R. Beltran, J. Joven, J. Camps. Immunohistochemical analysis of paraoxonases-1, 2, and 3 expression in normal mouse tissues. Free Radic. Biol. Med, 2008; 45(2), 146–157.

28. Devarajan A, Bourquard N, Hama S, Navab M, Grijalva VR, Morvardi S, Clarke CF, Vergnes L, Reue K, Teiber JF, Reddy ST. Paraoxonase 2 deficiency alters mitochondrial function and exacerbates the development of atherosclerosis. Antioxidants & Redox Signaling, 2011; 14(3), 341–351.

29. Giordano G, Cole TB, Furlong C E, Costa LG. Paraoxonase 2 (PON2) in the mouse central nervous system: a neuroprotective role? Toxicology and applied pharmacology, 2011; 256(3), 369–378.

30. Onaolapo AY, Ojo FO, Onaolapo OJ. Biflavonoid quercetin protects against cyclophosphamide-induced organ toxicities via modulation of inflammatory cytokines, brain neurotransmitters, and astrocyte immune reactivity. Journal of Pharmacology and Toxicology, 2023; 6(10), 1422.

31. Onaolapo AY, Adebisi EO, Adeleye AE, Olofinnade AT, Onaolapo OJ. Dietary Melatonin Protects Against Behavioural, Metabolic, Oxidative, and Organ Morphological Changes in Mice that are Fed High-Fat, High-Sugar Diet. Endocr Metab Immune Disord Drug Targets. 2020;20(4):570–583. doi: 10.2174/1871530319666191009161228.

32. Onaolapo AY, Adebisi EO, Adeleye AE, Olofinnade AT, Onaolapo OJ. Dietary Melatonin Protects Against Behavioural, Metabolic, Oxidative, and Organ Morphological Changes in Mice that are Fed High-Fat, High-Sugar Diet. Endocr Metab Immune Disord Drug Targets. 2020;20(4):570–583. doi: 10.2174/1871530319666191009161228.

33. Onaolapo OJ, Adeyemi OI, Amujoyegbe OJ, Fasola EA, Olofinnade AT, Onaolapo AY. High Dietary Fat Modulates Neurobehavioural Effect of Lopinavir/Ritonavir in Mice. Curr Pharm Biotechnol. 2020;21(2):158–168. doi: 10.2174/1389201020666191011144930.24.

34. Onaolapo OJ, Omotoso SA, Olofinnade AT, Onaolapo AY. Anti-Inflammatory, Anti-Oxidant, and Anti-Lipaemic Effects of Daily Dietary Coenzyme-Q10 Supplement in a Mouse Model of Metabolic Syndrome. Antiinflamm Antiallergy Agents Med Chem. 2021;20(4):380–388. doi: 10.2174/1871523020666210427111328

35. Olofinnade AT, Adeyeba A, Onaolapo AY, Onaolapo OJ. An assessment of the effects of azodicarbonamide-containing diet on neurobehaviour, brain antioxidant status and membrane lipid peroxidation status in rats. Central Nervous System Agents in Medicinal Chemistry (Formerly Current Medicinal Chemistry-Central Nervous System Agents), 2020; 20(1), 49–57.

36. Campos Alline C, Fogaca Manoela V, Aguiar Daniele C, Guimaraes Francisco S Animal models of anxiety disorders and stress. Revista Brasileira de Psiquiatria. 2013; 35: 101–111.

37. Olofinnade AT, Onaolapo AY, Onaolapo OJ. Olowe O.A. Hazelnut modulates neurobehaviour and ameliorates ageing-induced oxidative stress, and caspase-3-mediated apoptosis in mice. Current Aging Science, 2021; 14(2), 154–162.

38. Aziz SJ, Zeman-Pocrnich CE Tissue Processing. Methods in molecular biology (Clifton, N.J.), 2022; 2422, 47–63.

39. Smith, J. Aluminum Exposure in Human Populations. Environmental Health Journal, 2020; 10(2), 1–10. doi:10.1037/envh0000123

40. Bekhedda H, Menadi N, Demmouche A, Ghani A, Mai H. Histological study of the effects of aluminum chloride exposure on the brain of wistar rats female: the effects of aluminum chloride exposure on the brain of wistar rats female. J. Drug Delivery Ther. [Internet]. 2020;10(3-s):37–42.

41. Sheikh M, Ammar M. Efficacy of 5 and 10 mg donepezil in improving cognitive function in patients with dementia: a systematic review and meta-analysis. Frontiers in Neuroscience, 2024; 18, 1398952

42. Kadhim A, Ben Slima, A., Alneamah, G, Makni, M. Assessment of Histopathological Alterations and Oxidative Stress in the Liver and Kidney of Male Rats following Exposure to Aluminum Chloride. Journal of Toxicology, 2024;3997463.

43. Li Y, Yao J, Han C, Yang J, Chaudhry MT, Wang S, Liu H, Yin Y Quercetin, Inflammation and Immunity. Nutrients, 2016; 8:67. 10.3390/nu8030167

44. Chiang MC, Tsai TY, Wang CJ The Potential Benefits of Quercetin for Brain Health: A Review of Anti-Inflammatory and Neuroprotective Mechanisms. International journal of molecular sciences, 2023; 24(7), 6328. 10.3390/ijms24076328

45. Sarter M. Behavioural-cognitive targets for cholinergic enhancement. Current opinion in behavioral sciences, 2015;4, 22–26.

46. Sengoku R. Neuropathological features of cognitive decline in Lewy body dementia. Neurology and Clinical Neuroscience, 2020; 8(6), 356–361.

